# Regulatory properties of transcription factors with diverse mechanistic function

**DOI:** 10.1101/2023.06.15.545127

**Authors:** Md Zulfikar Ali, Sunil Guharajan, Vinuselvi Parisutham, Robert C. Brewster

## Abstract

Transcription factors (TFs) regulate the process of transcription through the modulation of different kinetic steps. Although models can often describe the observed transcriptional output of a measured gene, predicting a TFs role on a given promoter requires an understanding of how the TF alters each step of the transcription process. In this work, we use a simple model of transcription to assess the role of promoter identity, and the degree to which TFs alter binding of RNAP (stabilization) and initiation of transcription (acceleration) on three primary characteristics: the range of steady-state regulation, cell-to-cell variability in expression, and the dynamic response time of a regulated gene. We find that steady state regulation and the response time of a gene behave uniquely for TFs that regulate incoherently, i.e that speed up one step but slow the other. These TFs function as activators when regulating weak promoters but switch to repression when the promoters are strong or vice versa. Furthermore, we find that TFs with this regulatory make-up have dynamic implications, with one type of incoherent mode configuring the promoter to respond more slowly at intermediate TF concentrations. We also demonstrate that the noise of gene expression for these TFs is sensitive to promoter strength, with a distinct non-monotonic profile that is apparent under stronger promoters. Taken together, our work uncovers the coupling between promoters and TF regulatory modes with implications for understanding natural promoters and engineering synthetic gene circuits with desired expression properties.

## INTRODUCTION

Gene regulation is an essential process that controls a cell’s response to both external and internal stimuli. Accordingly, the ability to predict gene expression levels using forward modeling approaches has been a focus of in the field of regulatory biology [1–5]. These models have the potential to impact a wide range of applications, from biomedical—where deciphering the regulatory genome may clarify the influence of mutations—to synthetic biology, where constructing circuits with robust, predictable responses facilitates biological designs with industrial and therapeutic applications. However, achieving a model that is broadly useful has been a notoriously difficult problem. Recently, the need for precise quantitative models of gene input-output functions has been highlighted by studies demonstrating the sensitivity of essential regulatory networks to transcription factor (TF) dosage [6–9].

The regulation of a specific promoter hinges on a complex interplay of factors including the number and position of TF binding sites, the RNA polymerase (RNAP) recognition sequence, and the concentration and activity of TFs in a given cellular context. While numerous techniques allow for the measurement of the concentration and binding affinities of TFs, as well as TF occupancy across the genome, the combinatorial complexity intrinsic to promoters – regulated by various combinations of TFs binding at different locations – presents a formidable challenge to understanding the fundamental “rules” of gene regulation using natural promoters alone [10–13]. In contrast, highthroughput assays that measure the expression of thousands of synthetically designed promoters have yielded a more comprehensive understanding of systematic perturbations to promoters, testing the composability of characterized regulatory “parts” [4, 14–17].

The actual regulatory function of a transcription factor (TF) once it binds to the promoter is a crucial detail that is challenging to infer. Traditionally, TFs have been classified as “activators” or “repressors” based on their overall regulatory function on a specific promoter. However, it has become evident that these labels alone are insufficient for predicting the role of a TF on a different gene or in a different context. One reason for this is that the mechanics of regulation can affect various kinetic steps of the transcription cycle, as evidenced by studies [18–23].

A simplified two-step model describes transcription in bacteria [18, 24, 25]. The first step involves the binding of RNA polymerase (RNAP) to the promoter to form a closed complex, and the second step involves a series of conformational changes that result in an open complex followed by promoter escape – both steps considered irreversible in this context. More nuanced models that distinguish between open complex formation, promoter escape, and their reversibility exist [26, 27], but are not covered here.

Studies have demonstrated that TFs can influence distinct kinetic steps in the transcription process [27–34]. Certain TFs have been found to regulate transcription by affecting the ability of RNAP to occupy the promoter, achievable either through steric hindrance [2, 35] or by forming energetically favorable interactions with RNAP [36–38]. On the other hand, other TFs act on the kinetic steps of open complex formation and promoter escape [23, 29]. Furthermore, it has been shown that individual TFs may impact both rates [39, 40].

Recent studies have formalized a model for gene regulation that accommodates regulation on these two steps of the transcription process (Fig. 1A) [20, 40–42]. This model generalizes the regulation process across both steps, makes no assumptions about the net role of any TF, and takes into account that a TF can influence one or both kinetic steps of transcription. TF function is quantified through two parameters that measure the fold-change in the likelihood of RNAP occupancy at the promoter or the rate of initiation when TF is bound as compared to these processes when TF is not bound. The model refers to changes in RNAP occupancy at the promoter as stabilizing/destabilizing (parameterized as *β*—stabilizing if *β >* 1, destabilizing if *β <* 1), and changes to the rate of open complex formation as accelerating/decelerating (parameterized by *α*—accelerating if *α >* 1, decelerating if *α <* 1). In previous research, we utilized this model to measure the regulatory parameters of the *E. coli* TF, CpxR, in relation to binding position and found that *α* and *β* values, and thus the net regulatory role of the TF, are strongly dependent on the binding location [40].

**FIG. 1.**
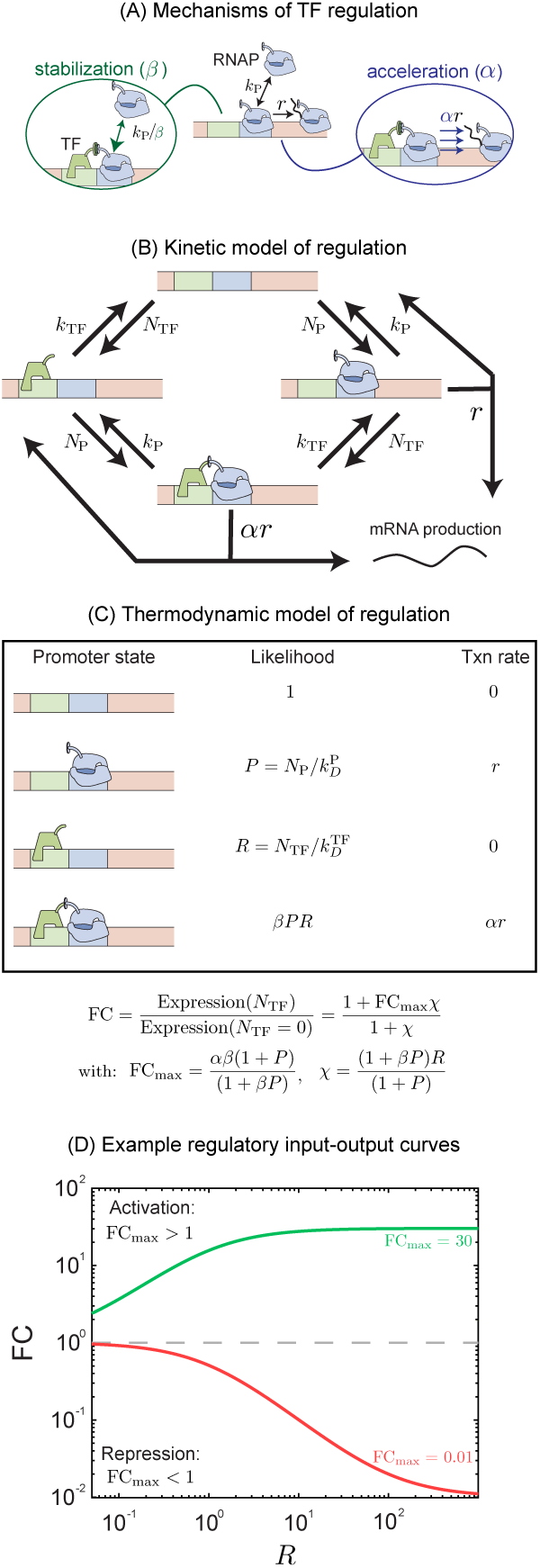
Schematic of regulation by TF in thermodynamic and kinetic model. (A) In this model the TF may function on two distinct steps of transcription related to the stabilization or destabilization of polymerase at the promoter and the acceleration or deceleration of transcription initiation by bound polymerase. (B) A kinetic model describing the transitions of the gene between 4 promoter states: empty, polymerase bound, TF bound and both TF and polymerase bound. Regulation is described by modifications to the rates of transition described here as *α* and *β*. (C) A simplified equilibrium model of the same system using the same parameters. (D) Example regulation curves for a TF which has an activating interaction FC_max_ *>* 1 (green curve) and for a TF which has a repressive interaction, FC_max_ *<* 1 (red curve).

Here we will consider the implications of this model using the simplest regulatory architecture, one where the promoter is regulated only by a single TF and that TF has only one binding site on the promoter. The possible regulatory outcomes of multi-TF regulation in this model are extremely broad but will not be discussed at length in this article. Here we use this model of gene regulation to explore the implications for regulation by a single TF in terms of three distinct qualities: the spectrum of responses a TF can have to different promoters at the mean expression level, the noise in expression, and the dynamic response time of a regulated gene. Furthermore, we investigate the intriguing phenomenon of autoregulation, wherein TFs regulate their own expression. Using thermodynamic and kinetic model approach we find that each of these properties depends sensitively on the proportion of each regulatory mechanism used by the regulating TF. Below we discuss the rich possibilities of gene regulatory responses and dynamics that result from the simple assumption that TFs are capable of regulating both the RNAP occupancy step and the transcription initiation steps of gene expression.

## RESULTS

### Model

The full kinetic model of gene regulation by a single TF is shown in Fig 1B. The model accounts for binding of TF and polymerase independently at rates that are proportional to the free TF (*N*_TF_) or polymerase concentration (*N*_P_). The TF and the polymerase unbind/dissociate from the bound states at the rate that is independent of the TF/polymerase concentration and is only dependent on the interaction between the TF and the polymerase, and binding site identity. The polymerase may also unbind through a productive initiation event where it will create an mRNA/protein. The regulatory role of the TF is encoded in two ways. The first, which we call stabilization, is represented as a constant factor, *β*, that alters the rate of TF and polymerase dissociation when co-bound. The second regulatory mechanism, acceleration *α*, is a constant multiplicative factor that modulates the initiation rate, *r*. These factors fall in the range between 0 and ∞; values greater than 1 represent regulatory interactions that promote gene expression (faster initiation or greater polymerase occupancy at the promoter), while values less than 1 represent regulatory interactions that repress expression (slower initiation or lower polymerase occupancy at the promoter). For simplicity and keeping the model tractable, we assume that the mRNA and protein production rates are incorporated in the initiation rate *r* and we do not include mRNA dynamics explicitly in the model.

First, we discuss the thermodynamic framework of the above-mentioned model where we compute the equilibrium probability of all the four possible states of gene expression namely, free, polymerase bound, TF bound, and co-bound states as shown in Fig. 1 [36, 43–48]. The advantage of this approach is that it tends to produce tractable analytic results at the cost of not explicitly considering if occupancy occurs through modulation of binding or unbinding rates of TFs or polymerases. The net fold-change predicted from this model is a function of the regulatory parameters, *α* and *β*, the number of TFs and polymerase present, *N*_TF_ and *N*_P_ as well as the affinity of the TF and polymerase for their binding sites, 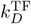 and 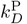. In this model we can write the fold-change, FC [40],

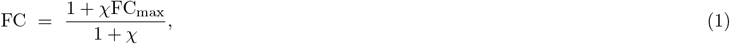

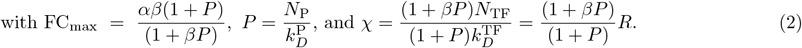

The fold-change curve can be neatly arranged into two parameters. First, *χ* is the effective concentration of the TF and depends on how many TFs are in the system compared to the equilibrium constant as well as a contribution from the stabilization regulatory interactions which can effectively increase (or decrease if *β <* 1). Second, FC_max_ is the fold-change of the gene when TFs are saturating; *i*.*e*. when *χ*≫1.

The model is greatly simplified under the assumption that the promoter is weak both without TF (*P*≪ 1) and with TF ((*βP*≪1); in this case, the parameters simplify greatly to 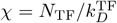 and FC_max_ = *αβ*. This result is intuitive, in this limit the maximum amount of regulation FC_max_ is the two mechanisms multiplied (how much more likely polymerase is to be at the promoter times how much faster it initiates transcription) and the effective number of TFs in the system *χ* is just the number of TFs divided by the equilibrium dissociation constant for the TF. In this limit, the two regulatory parameters and the two modes are not separable in their contribution to the average expression level of the gene. Example curves are shown in Fig. 1D that show stereotyped responses for repression (red) and activation (green) regulatory interactions. In all cases, the fold-change is monotonically increasing (activation) or decreasing (repression) until it approaches FC_max_.

### A phase space approach to visualizing regulatory function

One useful way to portray TF function is by plotting the regulatory function of a TF as a function of *α* and *β*. This is shown in Fig 2A with contour lines that mark FC_max_ in increments of 100−fold. The black contour represents FC_max_ = 1 which bifurcates the graph between activators (above the black line) and repressors (below the black line). The phase space is shown for two values of *P*, a weak promoter (left, *P* = 0.01) and a strong promoter (right, *P* = 1). When *P* is small, these lines simply follow the relationship *β* = FC_max_*/α* as indicated above which creates a straight line in log-log space corresponding to the weak limit described earlier. The lines only deviate from this relationship when *βP* is no longer significantly less than 1 or equivalently when *β*≫1. However, when *P* increases we see divergence from this simple relationship in the upper half of the plot where increasing *β* no longer increases the magnitude of the regulation (*i*.*e*. the contours become vertical).

**FIG. 2.**
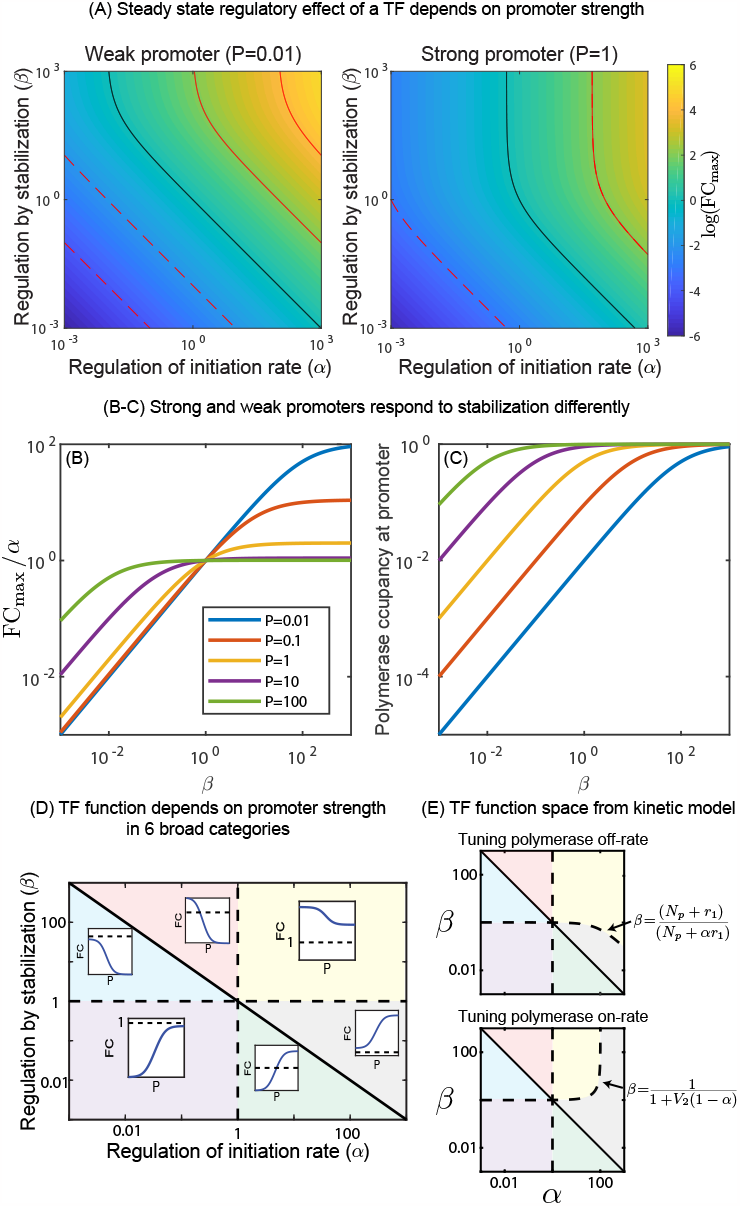
TF function predicted from regulatory parameters *α* and *β*. (A) The net “maximum” regulation from a TF as a function of the parameter *α* and *β* for a weak promoter (*P* = 0.01, left panel) and a strong promoter (*P* = 1, right panel). The observed regulation from a TF depends on promoter strength. (B-C) Positive stabilization is ineffective on strong promoters due to saturation of polymerase at the promoter. (D) The regulatory parameter space (*α*-*β* space) splits into 6 regions based on how a gene will respond to titration of promoter strength. The three lines that divide this space correspond to *αβ* = 1, *α* = 1 and *β* = 1. Insets in each region show the qualitative behavior of TFs from that region of parameter space as *P* is titrated. (E) The space is altered slightly if solved in the kinetic model; the boundary corresponding to *β* = 1 is now more complex depending on if the on-rate or off-rate of polymerase is tuned to change promoter strength. Here *V*_2_ is the ratio of the transcription initiation rate and the polymerase unbinding rate.

Fig. 2B shows contours of FC_max_*/α* for different values of *P* (left panel). As *β* becomes larger, FC_max_ simply becomes *αP/*(1 + *P*) and FC_max_ saturates as the occupancy of polymerase at the promoter goes to one (Fig. 2B, right panel). Furthermore, it can be seen that when *β >* 1 increasing *P* leads to a monotonic decrease in FC_max_, however when *β <* 1 we see the opposite, FC_max_ monotonically increases. This feature combined with the fact that phase-space of activation and repression varies with *P* generates an intricate pattern in phase-space.

#### Dependence of regulatory outcome on promoter strength

The interdependence of fold-change on regulatory parameters of TF (*α* and *β*) and promoter strength (*P*) divide the *α*−*β* space into a six regions by the lines *α* = 1, *β* = 1 and *αβ* = 1. This is diagrammed in Fig. 2D. Importantly, each region has a particular regulatory behavior when *P* is tuned independent of the TF concentration. The six regions respond to increasing values of *P* as follows

1. *αβ <* 1 and *β >* 1: low repression to high repression (blue region)
2. *αβ >* 1 and *α <* 1: activation to repression (red region)
3. *αβ <* 1 and *α >* 1: repression to activation (green region)
4. *αβ >* 1 and *β <* 1: low activation to high activation (gray region)
5. *α <* 1 and *β <* 1: high repression to low repression (purple region)
6. *α >* 1 and *β >* 1: high activation to low activation (yellow region)

It is important to note that this behavior is independent of the TF concentrations and solely depends on the TF regulatory parameters and the promoter strength. The most interesting behavior we observe is the switching of regulation from activation to repression in the red region, and repression to activation in the green region as *P* is tuned. The promoter strength at which the switch occurs is predicted to be *P*_*s*_ = (*αβ* −1)*/*(*β*−*αβ*). As a result, in the red shaded region, when *P < P*_*s*_ the TF behaves as an activator and when *P > P*_*s*_ as a repressor and vice versa for the green region. There is a simple qualitative way to understand this phenomenon. In the red region the regulatory interactions help stabilize polymerase at the promoter but also slow the transcription initiation process when polymerase is bound. For a weak promoter, this trade-off can result in a net increase in expression if *β* is greater than 1*/α* because the reduction in initiation is compensated by recruiting more polymerase at the promoter. For instance, as an example if *β* = 10 and *α* = 1*/*2, polymerase will occupy a weak promoter 10 times more which even if the TFs presence slows transcription by 1*/*2, this is a net activation. On the other hand, for strong promoters which may gain little or no additional polymerase occupancy from stabilizing interactions, the net change is that all transcription is slowed by 1*/*2 resulting in repression. It can also be seen from Eqn. 2 that when *P*→0 the FC_max_ becomes *αβ*. This means any reduction in fold-change due to reduction in initiation (*α <* 1) is compensated by an increase in the polymerase occupancy by stabilization. On the contrary when *P*→∞ the FC_max_ goes to *α* which is independent of the stabilization factor *β*. Consequently, *α <* 1 always leads to repression irrespective of the contribution from the stabilization. In both cases, we expect this phenomenon for TFs that are “incoherent” in their regulation; *i*.*e*. they slow one step while speeding up another. Specifically, it is predicted when the effect on *β* is the dominant interaction, in other words when the fold-stabilization is greater than the fold-deceleration or the fold-destabilization is greater than the fold-acceleration. Importantly, in previous studies, we have inferred that the *E. coli* TF CpxR regulates in this way at many binding positions [40].

#### Implications of the full kinetic model

The thermodynamic model outlined in Fig. 1C is agnostic to how stabilization *β* is incorporated in the model and also whether *P* is tuned through polymerase number affecting the net binding rate or the unbinding rate. To explicitly include these, and non-equilibrium effects, we turn to the full kinetic model outlined in Fig. 1B and solve for features such as the mean level of expression, the response time of single cells and the noise in gene expression using a chemical master equation approach (see SI). The expression for FC_max_ in the full model is more complex, given by

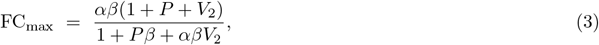

with a cumbersome expression of fold-change for finite TF concentrations (see SI). Here, *V*_2_ = *r/k*_P_ is the ratio of the transcription rate from polymerase bound state and the polymerase unbinding rate without TF (*k*_P,off_). In the limit *V*_2_→0 or alternatively when the transcription rate is much slower than the polymerase unbinding rate, the expression for FC_max_ reduces exactly to the thermodynamic model expression. The phase space of Fig. 2D changes in the full kinetic model. The phase-space is divided in the thermodynamic model by the lines *α* = 1, *β* = 1 and *αβ* = 1. In the full kinetic model, the boundaries of *α* = 1 and *αβ* = 1 remain unchanged, however, the third line changes depending on how *P* is tuned, *i*.*e*. by changing promoter affinity for polymerase *k*_P_ or by changing concentration of polymerase *k*_on,P_. Fig 2E shows both cases of these spaces. As can be seen, when *α*≤1 the lines between regions of space are unchanged (the red, blue and purple shaded regions). However, when *P* is tuned by changing the off-rate of polymerase from the promoter, the yellow region (activation becoming weaker) grows at the expense of the gray region (activation becoming stronger). However, the opposite is true when *P* is tuned by changing the on-rate of polymerase where the yellow region now shrinks at the expense of the gray region. The analytic expression for this line can be solved exactly and is shown next to each plot (also see Table I). All results that follow will be derived from the full kinetic model.

**TABLE 1.** Characterization of phase-space for FC_max_ from the thermodynamic and full kinetic model.

| Phase space for $FC_{\max}$ | | | | |
| --- | --- | --- | --- | --- |
| Model | Tuning $P$ through | $FC(R \rightarrow \infty, P \rightarrow 0)$ | $FC(R \rightarrow \infty, P \rightarrow \infty)$ | Line of separation |
| Thermodynamic | $k_P$ or $k_{\text{on},P}$ | $\alpha\beta$ | $\alpha$ | $\beta = 1$ |
| Kinetic model | $k_P$ | $\alpha\beta$ | $\frac{\alpha(N_P + r)}{N_P + \alpha r}$ | $\beta = (N_P + r)(N_P + \alpha r)^{-1}$ |
| | $k_{\text{on},P}$ | $\frac{\alpha\beta(1+V_2)}{1+\alpha\beta V_2}$ | $\alpha$ | $\beta = (1 + V_2(1 - \alpha))^{-1}$ |

#### Regulatory behaviors at finite TF concentration

To this point, we have discussed the dependence of the maximum fold-change possible from a TF, FC_max_, which will be observed when TF concentration is saturating. However, we also predict interesting phenomena at non-saturating TF concentrations. Most notably, although the maximum fold-change of a TF is a strictly increasing function with *α* and *β*, that is to say higher *α* and *β* always leads to an increase in the maximum fold-change observable from the regulated gene, at intermediate concentrations this is not necessarily the case. Figure 3A shows the parametric plot of fold-change, FC as a function of *α* and *β* but with a fixed, sub-saturating concentration of TF. Most notably, the top left corner of this plot which corresponds to low *α* and large *β* shows the minimum expected fold-change, even lower than expected for TFs whose regulatory role is both destabilizing and decelerating. Fig 3B shows contours of FC vs *β* at fixed *α*. As can be seen, below a certain value of *α*, higher stabilization actually decreases the fold-change. We can derive a simple relationship that determines if increased stabilization (*β*) will increase or decrease fold-change and the point of inflection between these two regimes occurs when *α* is a specific value *α*_*c*_ = *P/*(1 + *P* + *R*) (derived from the thermodynamic model. see SI Fig. S1 for comparison between the thermodynamic and full model). When TFs are saturating (*R*→∞) the inflection point occurs at *α* = 0 which is the allowable minimum for *α* and not realizable, and this is why we did not see this behavior in FC_max_ (Fig. 3A). However, when TF is at sub-saturating concentrations, this inflection point occurs at realizable levels of deceleration. Importantly, *α*_*c*_ is strictly decelerating, this phenomenon is not possible when the interaction is accelerating *α >* 1. Regardless, for a finite TF concentration (*R*) if *α* is less than *α*_*c*_ a TF with higher stabilization (*β*) produces a lower fold-change than a TF with lower stabilization. On the contrary, if *α* is greater than *α*_*c*_ then higher stabilization values will lead to a higher fold-change (see Fig. 3C).

**FIG. 3.**
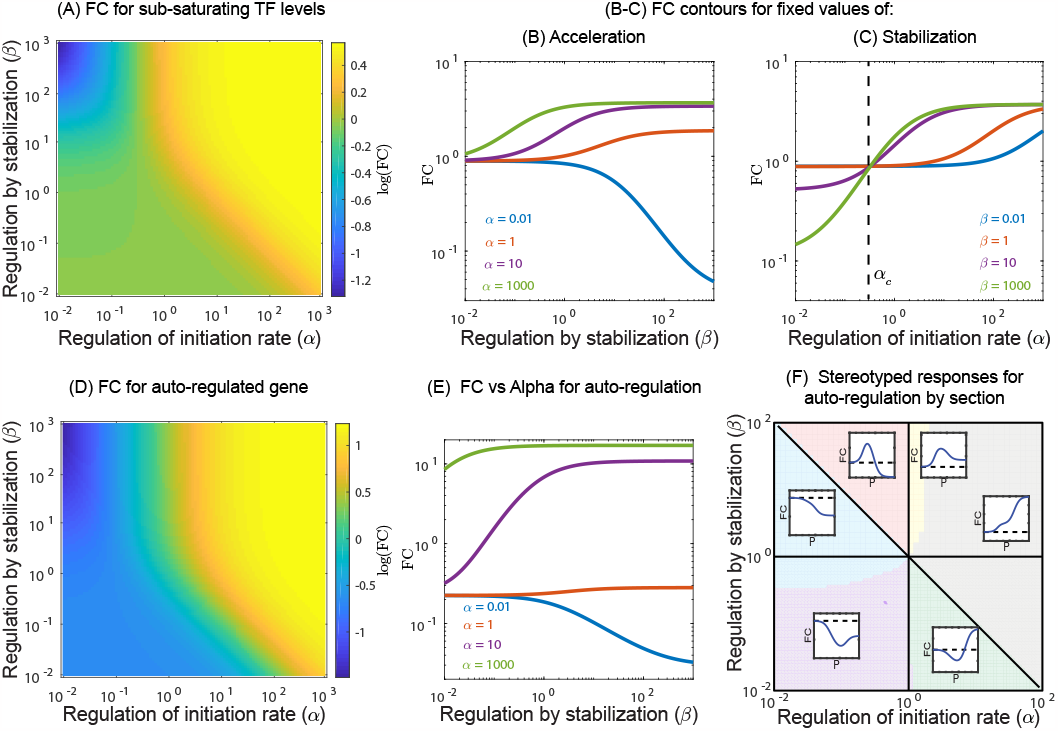
Regulation at non-saturating levels of TF. (A) Fold-change in expression as a function of *α* and *β* at a subsaturating TF concentration, *R* = 0.2. Here we see a curious feature where the upper left part of the phase space has the lowest fold-change. (B-C) Increasing the stabilization, *β*, increases fold-change for values of *α* above a critical value *α*_*c*_, however, when *α < α*_*c*_, the fold-change decreases with increased stabilization. The critical value, *α*_*c*_, can be exactly solved for and depends on *P* and *R*. (D-E) This phenomenon persists when examining autoregulatory genes. (F) Although the boundaries between regions of regulation space change for autoregulation, the same qualitative regions are seen, with the primary difference that *FC →* 1 as *P →* 0.

#### Regulatory behaviors of auto-regulation

We also consider the same simple regulatory architecture, a gene regulated only by one TF, but in the case where the gene itself produces the TF, *i*.*e*., an autoregulated gene. In this case, *N*_TF_ is no longer a parameter that we control but rather it is determined by the characteristics of the gene, the affinity of the TF for the binding site, the strength of the promoter *P* and the parameters of the regulatory interactions, *α* and *β*. The phase space of FC as a function of *α* and *β*, shown in Fig. 3D looks similar to that for finite TF number shown in panel A of the same figure. Although there are some quantitative differences. Interestingly, the same phenomenon occurs in autoregulation where a critical value of stabilization *α*_*c*_ exists, below which increased stabilization leads to lower fold-changes (Fig. 3E) Although, the analytic value of *α*_*c*_ is no longer the same and we do not have a simplified expression as in the case of simple regulation, we estimate *α*_*c*_ numerically and compare it with the full model for various promoter strength (see SI Fig. S1). Finally, we can again divide the parametric space of *α* and *β* into regions with qualitatively different responses to changing promoter strength *P* (Fig. 3F). Here we show that the same six regions exist that we observed previously for non-autoregulation (Fig. 2C). However, the line *β* = 1 no longer divides the regions of behavior, the dividing line severely reduces the size of the yellow region (strong activation that gets weaker with increased *P*) and moderately cuts into the purple region (strong repression that gets weaker with increased *P*).

### Response time

Next, we explore how the response time of a regulated gene is influenced by acceleration and stabilization. In order to compute response time, we assume that at time *t* = 0 the gene is expressing constitutively and at that time, TFs are instantly switched to an active state. Once actively regulated by TFs, the expression level will change before eventually reaching a new steady state. The response time is then computed as the time for the expression to reach halfway from the prior, unregulated state level to the new regulated level (see SI for details). We find that when *R* goes to infinity, the response times do not depend on the regulatory parameters *α* and *β*; the response time is one-cell cycle (SI Fig. S2B). When *R* approaches zero, the new steady state also approaches to the constitutive level and hence we cannot determine the response time. However, for intermediate values of *R*, the response time depends on the properties of the regulating TF. As an example, Fig. 4A shows three curves of expression as a function of time for different values of *α* and *β* having the same steady state fold-change 1.5. We find that for the same fold-change faster response time can be achieved by a TF having large *α*(425) and small *β*(0.01) (blue curve). For the same fold-change with *α* = 0.7 and *β* = 1000, the response time becomes slower by over one cell-cycle (yellow curve). A heat map showing the response time in *α−β* space is shown in Fig. 4B. We find that for strong stabilization and deacceleration (*α <* 1) the response times are the longest and reach several cell-cycles. A useful way to compare the response time is to examine response times of regulation as a function of fold-change. In Fig. 4C, we plot response time as a function of fold-change tuned by changing the stabilization, *β* at fixed *α* (top panel) or by changing acceleration *α* at fixed stabilization *β*. In the bottom panel of Fig. 4C, we see that for the same mean fold-change, TFs with lower *β* values (less stabilizing) have lower response times. In particular, depending on the value of *β* the response time decreases or increases monotonically with the fold-change. When the TF is a stabilizer (*β >* 1), increasing the fold-change decreases the response time. On the other hand, when the TF destabilizes (*β <* 1) increasing the fold-change increases the response time. Importantly, when *β* = 1 the response time is independent of the fold-change and is set by the constitutive gene (brown curve). In Fig. 3C we have shown that the fold-change is always monotonically increasing with *α*. This implies that when the TF is a stabilizer increasing *α* will lead to higher fold-change with faster response times. On the contrary, when the TF is a destabilizer increasing *α* will lead to higher fold-change with slower response times (4C).

**FIG. 4.**
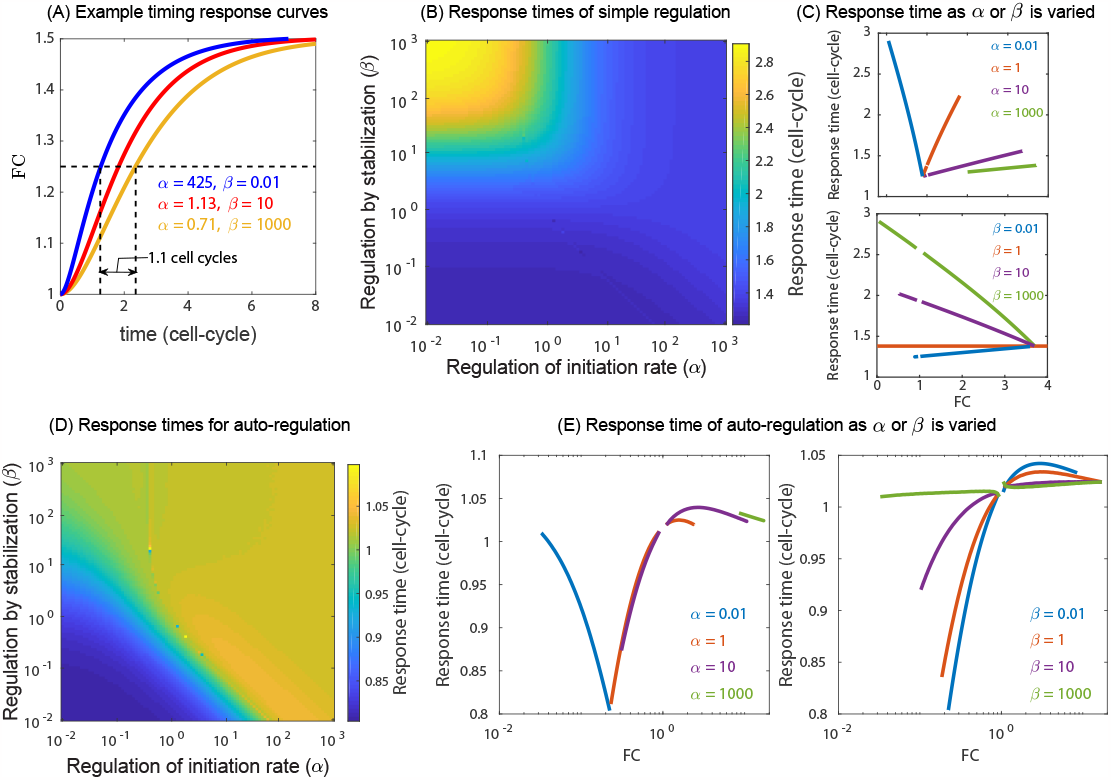
Response time of gene regulation depends on regulatory parameters. (A) Example traces of three TFs that have the same mean level of expression (FC) of a regulated gene at steady state. The time it takes to reach 50% of steady state varies substantially depending on the regulatory mechanism used by the TF. (B) Response time as a function of the parameters *α* and *β*. (C) Response time as a function of FC for TFs with a fixed *β* or *α* as the other parameter is varied. (D) Response times in the case of auto-regulation are significantly less variable (see scale). (E) Response time of auto-regulatory genes as a function of *FC* for fixed *α* or *β* while the other parameter is varied.

In the top panel of Fig. 4C we examine the response time when *α* is fixed and *β* is tuned to explore the response time as a function of fold-change. From Fig. 3B, we know that when *α* is below a critical value, increasing *β* will cause a decrease in fold-change. However, when *α* is above this threshold, increasing *β* will cause fold-change to increase. This is seen in the top panel of Fig. 4C, however, regardless of the specific value of *α*, the response time increases with *β*. This also implies that when TF is a repressor increasing fold-change reduces the response time (blue curve) and when the TF is an activator increasing fold-change increases the response time (green, purple, and brown curves).

We also examine how the response time depends on the regulatory parameters for an autoregulated gene. For an auto-regulated gene the number of TFs is a function of the regulatory parameters and hence the number of TFs are not fixed as we probe different *α* and *β*. As before, we assume that at time *t* = 0 the gene is expressing constitutively and at that time, TFs are instantly switched to an active state, i.e., the protein product of the gene binds to its own DNA and start regulating the gene. A heat map showing the response time in *α*−*β* space is shown in Fig. 4D. We find that for weak stabilization and weak acceleration the response times are the fastest and are lower than a cell-cycles. In fact, for the parameter regime explored we find that the response times hardly exceeds one cell cycle irrespective of the TF identity, an activator or a repressor. We also find an intermediate maximum in the response time as a function of both stabilization and acceleration which can be viewed in response time versus fold-change plots (Fig. 4E).

### Noise in expression

We compute noise as the coefficient of variation (CV) of the expression. In general, we will consider the fold-change in CV, which compares the change in noise from regulation to that of the unregulated (*R* = 0) case. For finite TF concentration the expression for CV is intractable and we solve it numerically (see SI). However when *R*→∞ the expression for noise (CV_max_) reduces to a simple form and is given by

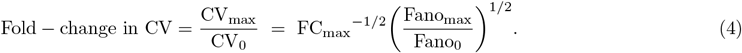

where, the fano-factor is

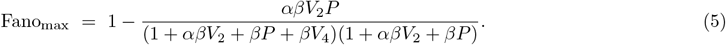

In the above expressions, setting *α* = *β* = 1 gives the noise of a constitutive gene (CV_0_ and Fano_0_). In Fig. 5, we examine how the noise is modulated as a function of TF concentration for various regulatory parameters and promoter strengths. For a pair of (*α, β*) and promoter strength (*P*) we vary *k*_TF,on_ as proxy for TF number in a range from (0, ∞) and examine the dependence of noise on the TF concentrations. When the number of TF is infinite the fold-change in CV approximately follows inverse square root of fold-change in expression shown as black line in Fig. 5B. In Fig. 5A, the solid lines show the trajectory of a typical repressing TF interactions as the number of TFs is titrated, while the dashed lines show activating interactions as the number of TFs is titrated for various promoter strengths and regulatory parameter *α, β*. We find three typical behaviors of the relationship between FC in CV and TF copy number. The first, colored blue, are cases where FC in CV is monotonically increasing (for repressing interactions) or decreasing (for activating interactions). The second type, colored red, show an intermediate maximum in FC in CV at a finite concentration of TFs. The last color, magenta, marks relationship where the FC in CV has both a maximum and a minimum at intermediate concentration. In some cases, the intermediate peak is a global maximum while in other cases the value at TF→∞ is greater than the intermediate peak. The occurrence of these relationships is shown in Fig. 5C-F for 4 different promoter strengths. We find that for very weak promoter the noise is always monotonic irrespective of whether it is an activator or a repressor (see Fig. 5C). For such promoters the noise is always higher than the constitutive gene if the TF is a repressor. On the contrary, the noise is always lower than the constitutive gene if the TF is an activator, i.e., the noise decreases monotonically with TF concentration and saturates at CV_max_. As we increase the promoter strength, the activators show a non-monotonic behavior (shown as red points in Fig. 5D-F) with an intermediate peak in the upper right corner of the phase-space which is greater than the noise of a constitutive gene. The regulatory parameters in the vicinity of *αβ* = 1 still show a monotonic noise. For a strong promoter, we find that the activators always show a non-monotonic behavior with an intermediate peak. However, the regulatory parameters corresponding to repressors show 3 different regimes (Fig. 5F); low *α* and *β* regime with monotonic noise (blue), a non-monotonic regime with a maximum (red), and a regime sandwiched in between where both a maximum and a minimum in noise is observed (magenta).

**FIG. 5.**
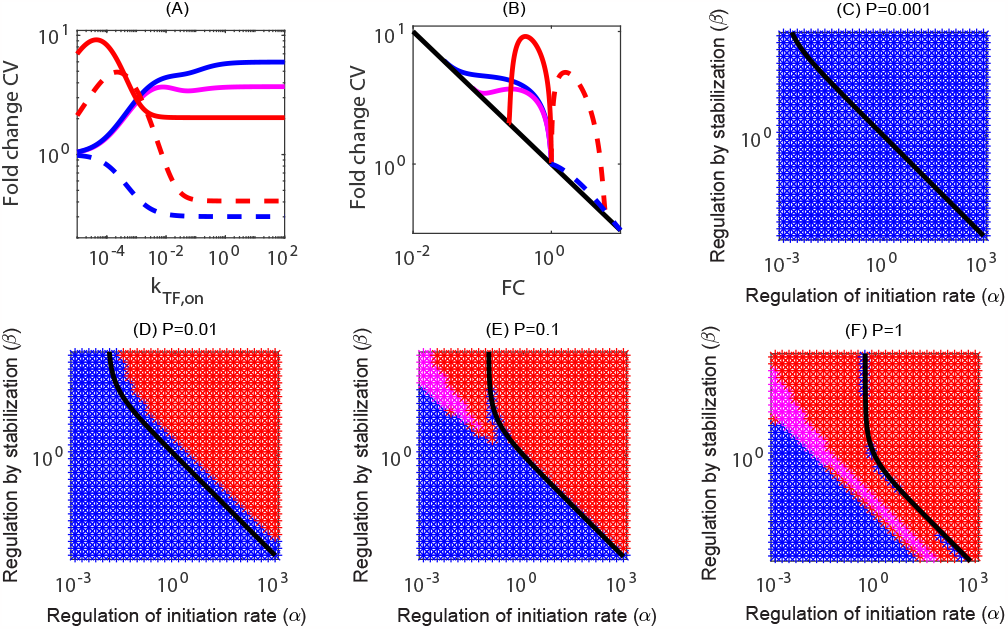
Phase-space showing monotonic and non-monotonic noise in gene expression as a function of TF concentration for various regulatory parameters. (A) The fold-change in CV as a function of TF concentration, *k*_TF,on_ can be monotonic (blue) (*i*.*e*. noise is highest for saturated TF concentrations), non-monotonic with a maximum at finite TF concentration (red), or non-monotonic with an internal maximum and minimum (magenta). The shape of this curve depends on both the promoter strength (*P*) and the regulatory parameters (*α, β*) of the TF. (C-F) The phase-space of noise for promoters of various strength. The color of the space in these plots specifies the qualitative shape of the fold-change in CV vs *k*_TF,on_ plot. The black line separates repressive TFs (left of the line) and activating TFs (right of the line). (C) For weak promoters, the noise is always monotonic. (D-F) For stronger promoters we see that activating TFs are always non-monotonic when the TF is activating (points to the right of the black line). However, in the repression regime, the noise can be monotonic (blue), or non-monotonic with a maximum (red) or both maximum and a minimum (magenta).

## CONCLUSIONS AND DISCUSSIONS

Here we have considered a simple model of TF function that accounts for regulatory interactions on two steps in gene expression process: occupancy of RNAP at the promoter and initiation of the transcription process. The realization of this model is that the characteristics of even the simplest regulatory systems are sensitive to the interplay of the regulatory interactions and the relative strength of the promoter. Primarily this interplay arises because the relative “value” of stabilizing interactions to a promoter depends on how well the promoter is capable of recruiting polymerase in the absence of TF; stronger promoters are altered less by stabilization than weak promoters. Although it is not in our model, acceleration should have a similar limitation set by the speed of transcriptional elongation. Our model assumes nothing about the relationship between these characteristics, only that they exist. In practice it may be that these modes are correlated in some way, for instance perhaps the fundamental interactions that produce stabilizing interactions also tend to be decelerating. However, in a past study we found stabilizing TFs with both accelerating and decelerating interactions; the methodology in that study was unable to resolve destabilizing interactions [40].

Among the most striking consequences of this model is that the regulatory function of a TF depends strongly on the promoter it is regulating. It is common to classify TFs as “activators” or “repressors” and presume that this is a meaningful classification, however, the implications of this model is that the net regulatory action of a TF is a complex interplay of these two mechanisms and, as such, is not a good classifier of TFs. Even the broad, intuitive characterizations such as “more stabilizing TFs should produce larger fold-changes” fails in certain situations (see Fig. 3A). Recently, we have seen that regulation by CpxR appears to be both stabilizing (*β >* 1) and decelerating (*α <* 1) for many binding locations (blue and red region of Fig. 2C). TFs with such “incoherent” regulatory interactions are particularly prone to be difficult to predict even their qualitative regulatory features on a promoter based on measurements of a different promoter. Indeed regulation data collected from the regulation of natural targets of CpxR show a mix of activating and repressing interactions even when the binding site is in similar locations on different promoters. There are a plethora of contextual information that could explain this phenomenon (interactions with other TFs on the promoter, network or off-target effects, *etc*.), however, the results here show that this is an expected behavior even in the absence of those higher-order interactions.

Regulatory modes impact more than just the steady-state levels of expression. We show that the response time of a gene to regulation depends on the regulatory modes of the TF. For simple regulation, the heat map of response times almost resembles an “AND” logic gate where a combination of stabilization and deceleration will cause the response of the gene to be slow with the possibility of doubling the response time compared to faster combinations. We do not see this same phenomenon for auto-regulation where the space is considerably more confined, however in all cases auto-repressive circuits respond faster than auto-activation. As a general phenomenon, it is known from simple models that auto-repression speeds up responses while auto-activation slows response times [49, 50], we show here that based on the regulatory parameters of the TF, the response times can vary substantially, although the general phenomenon that auto-repression speeds up responses while auto-activation slows response times is broadly true in this case [50, 51].

We further investigated the impact of the regulatory modes in the noise defined as the coefficient of variation (CV) of gene expression. We find three characteristic feature of noise as a function of fold-change. For weak promoter the noise is always monotonically increasing with the fold-change irrespective of whether the TF is an activator or a repressor. On the other hand for strong promoters, the noise can be monotonically increasing or non-monotonic with a maximum at intermediate fold-change or both a maximum and a minimum depending on the regulatory parameters. Overall, the balance of two regulatory interactions significantly alters every property of a TFs function. Although there are infinitely many combinations of *α* and *β* to achieve a given average fold-change, specific values of these parameters will alter the response dynamics of the gene, the noise and even how that TF will regulate other promoters. In this case, the broad labels of “activator” or “repressor” will not capture anything general about a TFs interaction.

However, our knowledge of the mechanisms of action of TFs is relatively limited, an important next step is to understand what part of this phase space is actually seen in real TFs.

## Appendix A: Model

The full kinetic model of gene regulation by a single TF is shown in Fig 1B. For simplicity we assume that there is only one binding site for TF and one for the polymerase. The model accounts for binding of TF and polymerase independently at rates that are proportional to the free TF (*N*_TF_) or polymerase concentration (*N*_P_) and their corresponding binding rates *k*_on,TF_ and *k*_on,P_. The TF and the polymerase unbind/dissociate from the bound states at the rate that is independent of the TF/polymerase concentration and is only dependent on the interaction between the TF and the polymerase, and binding site identity. We denote the unbinding rates of the TF and polymerase as *k*_TF_ and *k*_P_, respectively. The polymerase may also unbind through a productive initiation event where it will create an mRNA/protein. The regulatory role of the TF is encoded in two ways. The first, which we call stabilization, is represented as a constant factor, *β*, that alters the rate of TF and polymerase dissociation when co-bound, i.e., the unbinding rates of TF and polymerase from the co-bound states are altered to *β*^*−*1^*k*_TF_ and *β*^*−*1^*k*_P_. The second regulatory mechanism, acceleration (*α*), is a constant multiplicative factor that modulates the initiation rate, *r*. These factors fall in the range between 0 and ; values greater than 1 represent regulatory interactions that promotes gene expression (faster initiation or greater polymerase occupancy at the promoter), while values less than 1 represent regulatory interactions that repress expression (slower initiation or lower polymerase occupancy at the promoter. For simplicity and keeping the model tractable, we assume that the mRNA and protein production rates are incorporated in the initiation rate *r* and we do not include mRNA dynamics explicitly in the model. Finally, the functional product of the gene *m* is diluted at the rate *γ* which primarily occurs through cell division.

The chemical master equations (CMEs) governing the dynamics of gene expression (*m*) can then be given by the following equations,

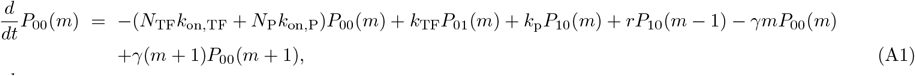

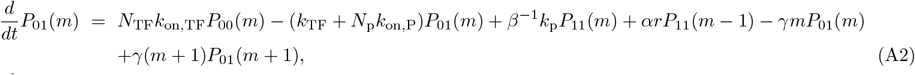

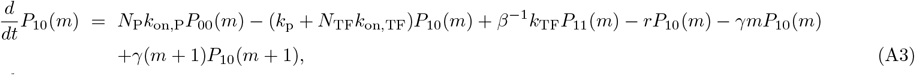

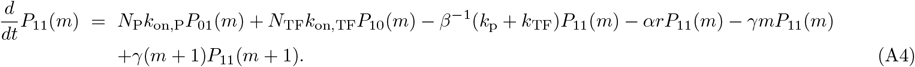

Here, *P*_*i,j*_(*m*) denotes the probability of having *m* proteins at time *t* in the state (*i, j*). *i* and *j* denotes the occupancy of polymerase and TF respectively and 0(or 1) indicates if the binding site is occupied (or free). Summing both side of the equations from *m* = 0 to *m* = ∞, we obtain the rate equation for mean occupancy in each state

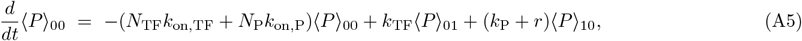

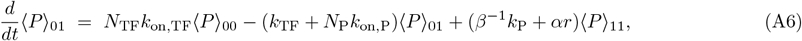

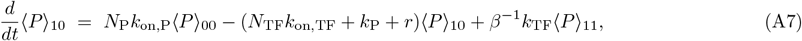

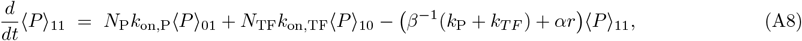

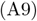

Similarly, multiplying both side of Eqns. A(1-4) by *m* and summing over *m* = 0 to *m* = ∞ give

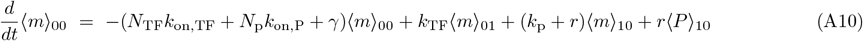

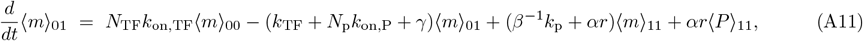

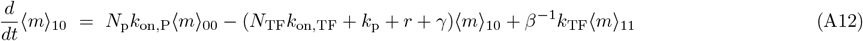

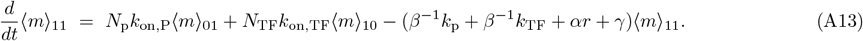

We obtain the equation for time evolution of the mean protein number (⟨*m*⟩) by adding Eqns. A(10-13)

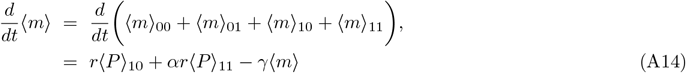

The steady state solutions can be obtained by setting the right hand side of the equations to zero which gives

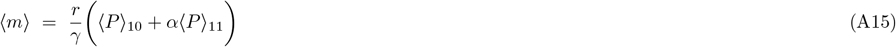

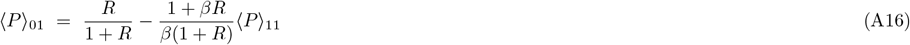

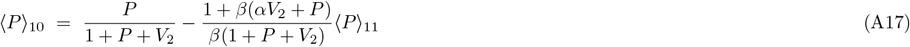

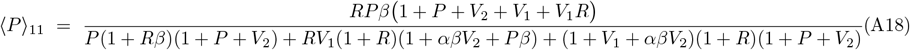

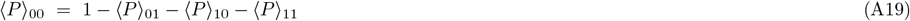

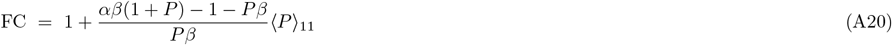

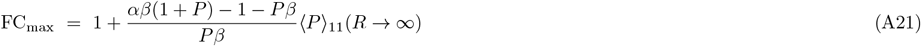

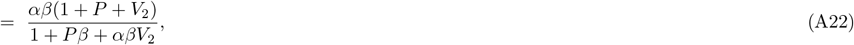

In the above equations we have substituted *R* = *N*_TF_*k*_on,TF_*/k*_TF_, *P* = *N*_P_*k*_on,P_*/k*_P_, *V*_1_ = *k*_TF_*/k*_P_, *V*_2_ = *r/k*_P_. Since ⟨*P*⟩ _11_≥0, for any concentration of TF, the nature of regulation (activation or repression) is determined by the expression *αβ*(1 + *P*)−1−*Pβ*. When this quantity is greater(less) than one we obtain activation(repression). Also the condition for fold-change of 1 is independent of TF concentration and is given by the equation

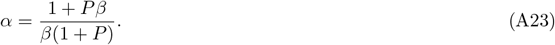

The polymerase unbinding rate we use is *k*_P_ = 1 s^*−*1^. The net polymerase binding rate (*N*_P_*k*_on,P_) is then determined from the promoter strength *P* = *N*_P_*k*_on,P_*/k*_P_ which is varied in the range 0.01−1, the weak promoter limit being 0.01 and 1 the strong promoter limit. Note that we do not tune the polymerase number independently. For the TF unbinding rate we use *k*_TF_ = 0.001s^*−*1^ corresponding to LacI binding site O1 in *E. Coli* [13]. We condense the TF number (*N*_TF_)and the sinlge TF binding rate (*k*_on,TF_) into one parameter which is varied in the range 0.0001−1. We use cell-division time (*τ*) of 40 minutes corresponding to *γ* = *ln*(2)*/τ* = 0.0003 s^*−*1^.

## Appendix B

Noise in gene expression

We compute coefficient of variation (CV) as a measure of noise. In order to do that we obtain the second moments by multiplying the Eqns. A(1-4) by *m*^2^ and summing over *m* = 0 to *m* = ∞ given by

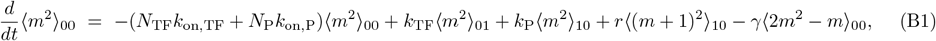

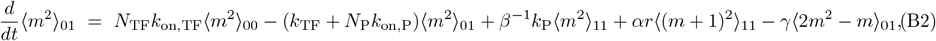

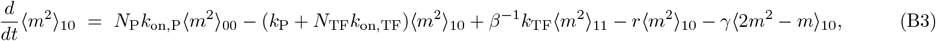

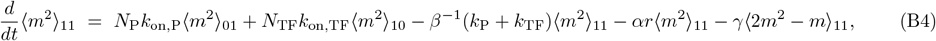

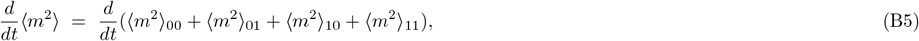

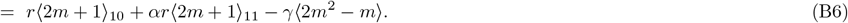

The steady state second moments are obtained by setting the above equations to zero which along with the steady states first moment gives

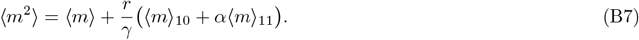

In order to compute CV or Fano factor we first numerically estimate the moments ⟨*m*⟩ _10_ and ⟨*m*⟩ _11_ and then use the the following equations,

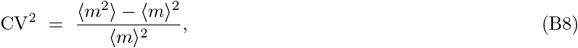

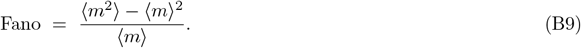

## Appendix C

Autoregulating gene

The set of ordinary differential equations governing the dynamics of an auto-regulating gene can be written as

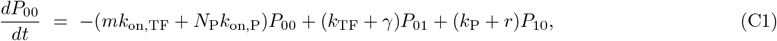

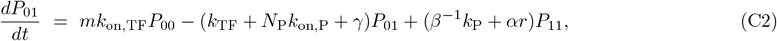

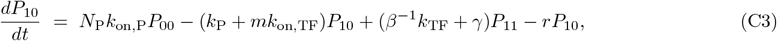

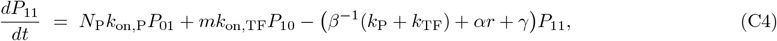

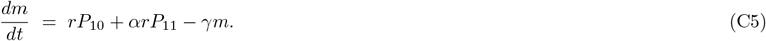

We numerically evaluate the steady state expression level by setting the above equations to zero. To compute the response time we solve the ordinary differential equation using ode45 in MATLAB.

## Appendix D

Response time

To compute the response time we solve the ordinary differential equations A(1-4) and A14 in MATLAB using the solver ode45. We assume that at time *t* = 0 the gene is expressing constitutively which gives the steady state expression level and the occupancies at *t* = 0 as

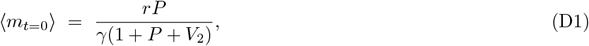

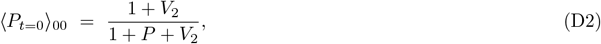

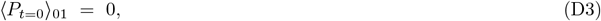

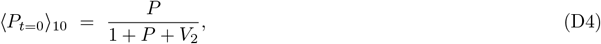

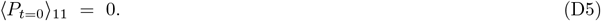

At time (*t* = 0), TFs are instantly switched to an active state. Once actively regulated by TFs, the expression level will change before eventually reaching a new steady state given by Eqns A(15-19). The response time is then computed as the time for the expression to reach halfway from the prior, unregulated state level to the new regulated level.

### SUPPLEMENTARY FIGURES

**FIG. S1.**
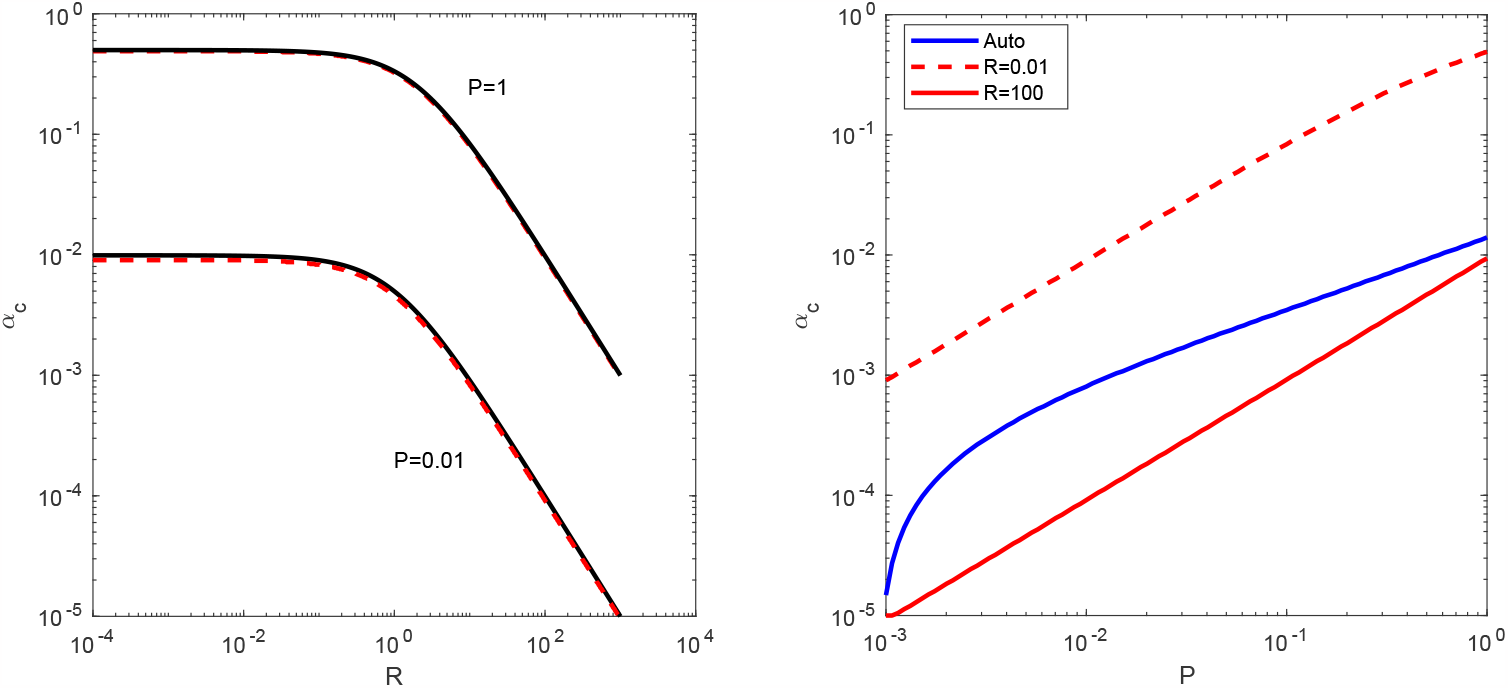
Critical acceleration, *α*_*c*_, for different models of gene regulation. (A) *α*_*c*_ from the thermodynamic model (black lines) and the full model (red lines) as a function of TF concentration for weak (*P* = 0.01) and strong promoter (*P* = 1). (B) *α*_*c*_ for autoregulated gene (blue line) and the genes regulated by fixed TF concentration (red lines) as a function of promoter strength.

**FIG. S2.**
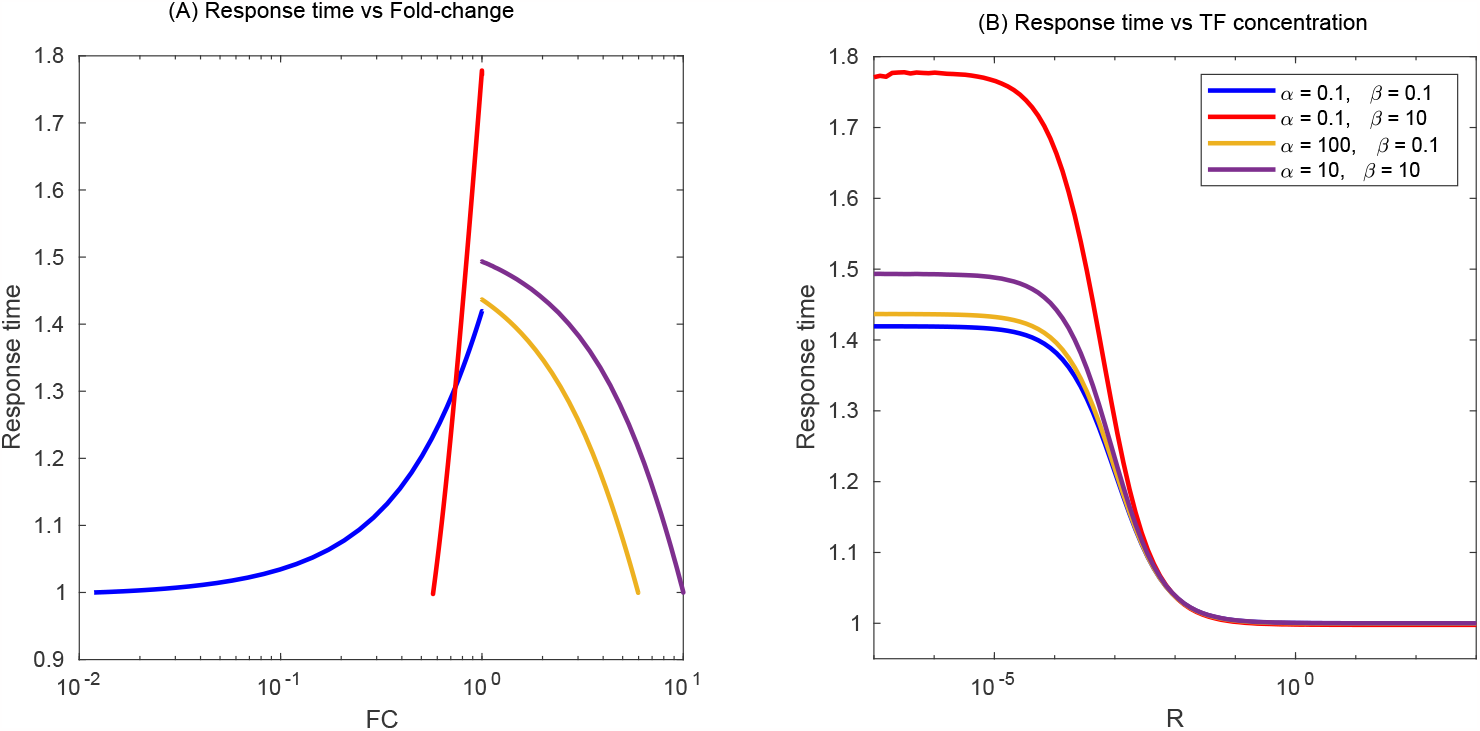
Response time when TF concentration is tuned. Response time as a function of fold-change (A) or TF concentraion (B) as the TF concentration is tuned. As the TF concentration approaches to infinity the response time is saturated to one cell-cycle irrespective of the regulatory parameter of the TF.

## Notes

### Competing Interest Statement

The authors have declared no competing interest.

## References

[1] J. Gunawardena, Time-scale separation–Michaelis and Menten’s old idea, still bearing fruit, FEBS J 281, 473 (2014).

[2] J. Gertz, E. D. Siggia, and B. A. Cohen, Analysis of combinatorial cis-regulation in synthetic and genomic promoters, Nature 457, 215 (2009).

[3] F. Spitz and E. E. Furlong, Transcription factors: from enhancer binding to developmental control, Nat Rev Genet 13, 613 (2012).

[4] M. Levo and E. Segal, In pursuit of design principles of regulatory sequences, Nat Rev Genet 15, 453 (2014).

[5] M. A. White, J. C. Kwasnieski, C. A. Myers, S. Q. Shen, J. C. Corbo, and B. A. Cohen, A Simple Grammar Defines Activating and Repressing cis-Regulatory Elements in Photoreceptors, Cell Rep 17, 1247 (2016).

[6] S. Naqvi, S. Kim, H. Hoskens, H. S. Matthews, R. A. Spritz, O. D. Klein, B. Hallgrímsson, T. Swigut, P. Claes, J. K. Pritchard, and J. Wysocka, Precise modulation of transcription factor levels reveals drivers of dosage sensitivity, bioRxiv 10.1101/2022.06.13.495964 (2022).

[7] S. Uphoff, N. D. Lord, B. Okumus, L. Potvin-Trottier, D. J. Sherratt, and J. Paulsson, Stochastic activation of a DNA damage response causes cell-to-cell mutation rate variation, Science 351, 1094 (2016).

[8] A. F. Johnson, H. T. Nguyen, and R. A. Veitia, Causes and effects of haploinsufficiency, Biol Rev Camb Philos Soc 94, 1774 (2019).

[9] R. Zug, Developmental disorders caused by haploinsufficiency of transcriptional regulators: a perspective based on cell fate determination, Biol Open 11 (2022).

[10] N. J. Guido, X. Wang, D. Adalsteinsson, D. McMillen, J. Hasty, C. R. Cantor, T. C. Elston, and J. J. Collins, A bottom-up approach to gene regulation, Nature 439, 856 (2006).

[11] D. Sprinzak and M. B. Elowitz, Reconstruction of genetic circuits, Nature 438, 443 (2005).

[12] M. R. Atkinson, M. A. Savageau, J. T. Myers, and A. J. Ninfa, Development of genetic circuitry exhibiting toggle switch or oscillatory behavior in Escherichia coli, Cell 113, 597 (2003).

[13] M. Z. Ali, V. Parisutham, S. Choubey, and R. C. Brewster, Inherent regulatory asymmetry emanating from network architecture in a prevalent autoregulatory motif, Elife 9, e56517 (2020).

[14] S. Kosuri, D. B. Goodman, G. Cambray, V. K. Mutalik, Y. Gao, A. P. Arkin, D. Endy, and G. M. Church, Composability of regulatory sequences controlling transcription and translation in Escherichia coli, Proc Natl Acad Sci U S A 110, 14024 (2013).

[15] T. C. Yu, W. L. Liu, M. S. Brinck, J. E. Davis, J. Shek, G. Bower, T. Einav, K. D. Insigne, R. Phillips, S. Kosuri, and G. Urtecho, Multiplexed characterization of rationally designed promoter architectures deconstructs combinatorial logic for IPTG-inducible systems, Nat Commun 12, 325 (2021).

[16] E. Sharon, D. van Dijk, Y. Kalma, L. Keren, O. Manor, Z. Yakhini, and E. Segal, Probing the effect of promoters on noise in gene expression using thousands of designed sequences, Genome Res 24, 1698 (2014).

[17] D. van Dijk, E. Sharon, M. Lotan-Pompan, A. Weinberger, E. Segal, and L. B. Carey, Large-scale mapping of gene regulatory logic reveals context-dependent repression by transcriptional activators, Genome Res 27, 87 (2017).

[18] W. R. McClure, Mechanism and control of transcription initiation in prokaryotes, Annu Rev Biochem 54, 171 (1985).

[19] L. J. Friedman and J. Gelles, Mechanism of transcription initiation at an activator-dependent promoter defined by singlemolecule observation, Cell 148, 679 (2012).

[20] C. Scholes, A. H. DePace, and nchez, Combinatorial Gene Regulation through Kinetic Control of the Transcription Cycle, Cell Syst 4, 97 (2017).

[21] N. J. Fuda, M. B. Ardehali, and J. T. Lis, Defining mechanisms that regulate RNA polymerase II transcription in vivo, Nature 461, 186 (2009).

[22] D. J. Lee, S. D. Minchin, and S. J. Busby, Activating transcription in bacteria, Annu Rev Microbiol 66, 125 (2012).

[23] D. Jensen, A. R. Manzano, J. Rammohan, C. L. Stallings, and E. A. Galburt, CarD and RbpA modify the kinetics of initial transcription and slow promoter escape of the Mycobacterium tuberculosis RNA polymerase, Nucleic Acids Res 47, 6685 (2019).

[24] G. N. Gussin, Kinetic analysis of RNA polymerase-promoter interactions, Methods Enzymol 273, 45 (1996).

[25] M. Morrison, M. Razo-Mejia, and R. Phillips, Reconciling kinetic and thermodynamic models of bacterial transcription, PLoS Comput Biol 17, e1008572 (2021).

[26] W. R. McClure, Rate-limiting steps in RNA chain initiation, Proc Natl Acad Sci U S A 77, 5634 (1980).

[27] D. Jensen and E. A. Galburt, The Context-Dependent Influence of Promoter Sequence Motifs on Transcription Initiation Kinetics and Regulation, J Bacteriol 203 (2021).

[28] F. Rojo, Repression of transcription initiation in bacteria, J Bacteriol 181, 2987 (1999).

[29] J. Xu and G. B. Koudelka, Repression of transcription initiation at 434 P(R) by 434 repressor: effects on transition of a closed to an open promoter complex, J Mol Biol 309, 573 (2001).

[30] G. Lloyd, P. Landini, and S. Busby, Activation and repression of transcription initiation in bacteria, Essays Biochem 37, 17 (2001).

[31] A. Barnard, A. Wolfe, and S. Busby, Regulation at complex bacterial promoters: how bacteria use different promoter organizations to produce different regulatory outcomes, Curr Opin Microbiol 7, 102 (2004).

[32] D. F. Browning and S. J. Busby, The regulation of bacterial transcription initiation, Nat Rev Microbiol 2, 57 (2004).

[33] S. A. van Hijum, M. H. Medema, and O. P. Kuipers, Mechanisms and evolution of control logic in prokaryotic transcriptional regulation, Microbiol Mol Biol Rev 73, 481 (2009).

[34] Y. Feng, Y. Zhang, and R. H. Ebright, Structural basis of transcription activation, Science 352, 1330 (2016).

[35] H. G. Garcia and R. Phillips, Quantitative dissection of the simple repression input-output function, Proc Natl Acad Sci U S A 108, 12173 (2011).

[36] T. Kuhlman, Z. Zhang, M. H. S. Jr., and T. Hwa, Combinatorial transcriptional control of the lactose operon of Escherichia coli, Proc Natl Acad Sci U S A 104, 6043 (2007).

[37] X. He, M. A. Samee, C. Blatti, and S. Sinha, Thermodynamics-based models of transcriptional regulation by enhancers: the roles of synergistic activation, cooperative binding and short-range repression, PLoS Comput Biol 6 (2010).

[38] J. B. Kinney, A. Murugan, J. C. G. Callan, and E. C. Cox, Using deep sequencing to characterize the biophysical mechanism of a transcriptional regulatory sequence, Proc Natl Acad Sci U S A 107, 9158 (2010).

[39] J. Blau, H. Xiao, S. McCracken, P. O’Hare, J. Greenblatt, and D. Bentley, Three functional classes of transcriptional activation domain, Mol Cell Biol 16, 2044 (1996).

[40] S. Guharajan, S. Chhabra, V. Parisutham, and R. C. Brewster, Quantifying the regulatory role of individual transcription factors in escherichia coli, Cell Reports 37, 109952 (2021).

[41] F. Wong and J. Gunawardena, Gene Regulation in and out of Equilibrium, Annu Rev Biophys 49, 199 (2020).

[42] R. Martinez-Corral, M. Park, K. Biette, D. Friedrich, C. Scholes, A. S. Khalil, J. Gunawardena, and A. H. DePace, Transcriptional kinetic synergy: a complex landscape revealed by integrating modelling and synthetic biology, bioRxiv 10.1101/2020.08.31.276261 (2020), https://www.biorxiv.org/content/early/2020/08/31/2020.08.31.276261.full.pdf.

[43] G. K. Ackers, A. D. Johnson, and M. A. Shea, Quantitative model for gene regulation by lambda phage repressor, Proc Natl Acad Sci U S A 79, 1129 (1982).

[44] L. Bintu, N. E. Buchler, H. G. Garcia, U. Gerland, T. Hwa, J. Kondev, and R. Phillips, Transcriptional regulation by the numbers: models, Curr Opin Genet Dev 15, 116 (2005).

[45] J. B. Kinney, A. Murugan, C. G. Callan, and E. C. Cox, Using deep sequencing to characterize the biophysical mechanism of a transcriptional regulatory sequence, Proc. Natl. Acad. Sci. U.S.A. 107, 9158 (2010).

[46] N. E. Buchler, U. Gerland, and T. Hwa, On schemes of combinatorial transcription logic, Proc Natl Acad Sci U S A 100, 5136 (2003).

[47] J. M. Vilar and S. Leibler, DNA looping and physical constraints on transcription regulation, J Mol Biol 331, 981 (2003).

[48] H. G. Garcia, A. Sanchez, J. Q. Boedicker, M. Osborne, J. Gelles, J. Kondev, and R. Phillips, Operator sequence alters gene expression independently of transcription factor occupancy in bacteria, Cell Rep 2, 150 (2012).

[49] N. Rosenfeld, M. B. Elowitz, and U. Alon, Negative autoregulation speeds the response times of transcription networks, J Mol Biol 323, 785 (2002).

[50] M. Z. Ali and R. C. Brewster, Controlling gene expression timing through gene regulatory architecture, PLoS Computational Biology 18, e1009745 (2022).

[51] A. D. Co, M. C. Lagomarsino, M. Caselle, and M. Osella, Stochastic timing in gene expression for simple regulatory strategies, Nucleic Acids Res 45, 1069 (2017).

